# Sample Preparation by Easy Extraction and Digestion (SPEED) - A Universal, Rapid, and Detergent-free Protocol for Proteomics based on Acid Extraction

**DOI:** 10.1101/393249

**Authors:** Joerg Doellinger, Andy Schneider, Marcell Hoeller, Peter Lasch

## Abstract

Mass spectrometry is the method of choice for deep and comprehensive analysis of proteomes and has become a key technology to support the progress in life science and biomedicine. However, sample preparation in proteomics is not standardized and contributes to a lack of reproducibility. The main challenge is to extract all proteins in a manner that enables efficient digestion into peptides and is compatible with subsequent mass spectrometric analysis. Current methods are based on the idea of removing detergents or chaotropic agents during sample processing, which are essential for protein extraction but interfere with digestion and LC-MS. These multi-step preparations are prone to losses, biases and contaminations, while being time-consuming and labor-intensive.

We report a universal detergent-free method, named *Sample Preparation by Easy Extraction and Digestion (SPEED)*, which is based on a simple three-step procedure, acidification, neutralization and digestion. SPEED is a one-pot method for peptide generation from various sources and is easily applicable even for lysis-resistant sample types as pure trifluoroacetic acid (TFA) is used for highly efficient protein extraction. SPEED-based sample processing is highly reproducible, provides exceptional peptide yields and enables preparation even of tissue samples with less than 15 min hands-on time and without any special equipment. Evaluation of SPEED performance revealed, that the number of quantified proteins and the quantitative reproducibility are superior compared to the well-established sample processing protocols FASP, ISD-Urea and SP3 for various sample types, including human cells, bacteria and tissue, even at low protein starting amounts.

## INTRODUCTION

Proteins regulate and catalyze all cellular processes. The analysis of the entity of proteins is therefore of utmost importance for a molecular understanding of life. During the last years, technical progress in the fields of mass spectrometry (MS) and bioinformatics has enabled the realization of deep and reproducible proteomic studies at increased throughput ^1^. The majority of these studies are currently performed using a bottom-up approach, which relies on the digestion of proteins into smaller peptides ^2^. Different sample preparation methods have been developed aiming at enabling comprehensive and reproducible generation of peptides from proteomes extracted from a large variety of sample types. Current protocols employ detergents, e.g. sodium dodecyl sulfate (SDS), or chaotropic agents, such as urea, for protein extraction and support sample lysis by physical disruption methods, such as heat, ultra-sonication or grinding ^3–5^. As many extraction reagents inhibit enzymatic digestion of proteins and are incompatible with LC-MS/MS, the idea behind most sample preparation methods is to remove interfering substances prior to digestion either by filtration (FASP) ^6^, precipitation (on-pellet digestion, STrap) ^7,8^ or bead-based purification (SP3) ^9^. However, enhancing lysis by use of physical disruption methods and subsequent protein purification requires additional sample handling steps, which are associated with attendant losses, biases and possible contaminations, while being time-consuming and labor-intensive.

The main exception is the well-established and frequently used in-solution digestion (ISD) of proteins, which is most frequently based on the extraction of proteins at high urea concentrations and subsequent dilution into a concentration range which is not inhibiting tryptic digestion anymore ^10^. Urea-based ISD (ISD-Urea) benefits greatly form the reduced number of individual steps needed for sample preparation and is known to be robust, highly reproducible and easy to perform. However, ISD-Urea is not a universal method for bottom-up proteomics as urea is in contrast to strong detergents, such as SDS, ineffective for extracting proteins from hard-to-lyse samples, such as tissues or gram-positive bacteria. Furthermore urea is known to attach artificial modifications to proteins, known as carbamylations ^11^. This prevents samples from being lysed at elevated temperatures, which would be beneficial for proteome extraction especially from challenging samples. The limitations of in-solution digestion arise from the compromise between efficient lysis and low interference with enzymatic activity of primarily trypsin.

This study presents the development of a sample preparation method, which overcomes the limitations in lysis and extraction of ISD but preserves its straight-forward approach. The new method is termed *Sample Preparation by Easy Extraction and Digestion (SPEED)* and consists of three simple steps, namely acidification, neutralization and digestion. SPEED uses neither detergents nor chaotropic agents for protein extraction and is fast, cheap, robust, highly-reproducible, well-suited for various sample types and easy to perform even for non-experts. The comparison of SPEED with FASP, SP3 and ISD-Urea for label-free quantitative analysis of eukaryotic cells and tissue as well as different bacterial species shows its universal applicability and superior performance especially concerning the number of quantified proteins and quantitative reproducibility over a wide range of sample types.

## RESULTS

### Development of SPEED

Comprehensive, accurate and deep proteome analysis relies on the reproducible generation of peptides from the entity of proteins of a given sample type. The main challenge during sample preparation is to extract all proteins in a manner that enables efficient digestion into peptides and is compatible with subsequent mass spectrometric analysis. A universal bottom-up proteomics sample preparation method would ideally consist of a single extraction step, which is independent of the sample type, and allows subsequent proteolytic digestion without the need for protein purification. In order to develop such a one-step universal protein extraction method, we decided to leave the track of common lysis buffer chemistry and ended up using pure trifluoroacetic acid (TFA).

TFA is a strong acid (pK_a_ = 0,2) and an excellent solvent for proteins ^12^. We found that pure TFA is able to dissolve cells and tissues within few minutes at room temperature forming clear lysates. Viscosity of the lysates is low as DNA is degraded rapidly after acidification. Analysis of *E.coli* cells incubated with TFA for different times between 1-60 min revealed, that TFA does neither hydrolyze peptide bonds nor modifies amino acid residues (Fig. 1). Protein intensities of samples with different TFA-incubation times had excellent Pearson coefficients of at least 0.994 and numbers of protein and peptide identifications were unaffected (Fig. 1).

**Figure 1:**
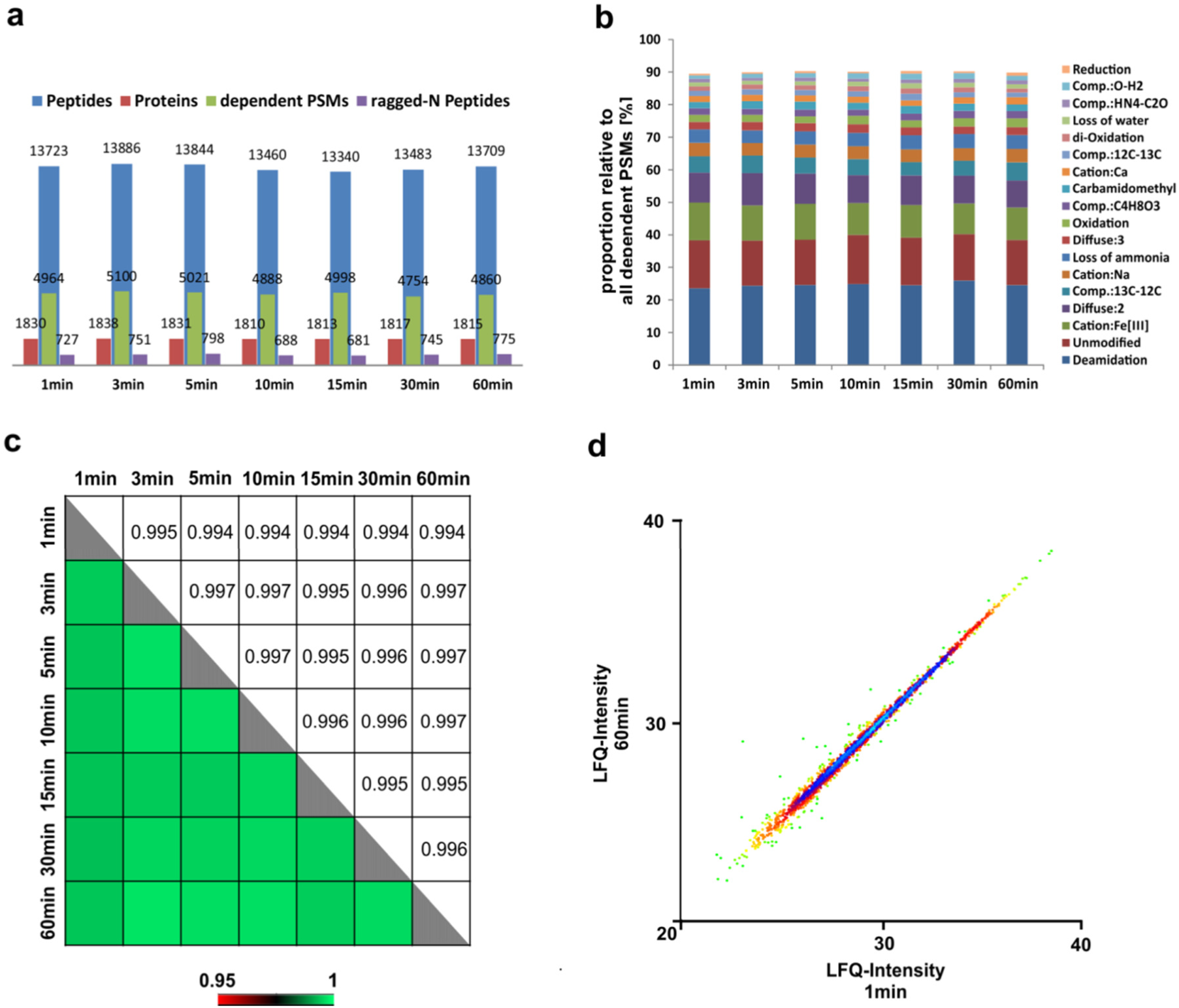
Effect of TFA incubation time. The effect of TFA incubation time ranging from 1 to 60 min on the performance of an *E. coli* proteome analysis is shown. The number of peptide and protein identifications as well as ragged-N peptides and the total number of peptide spectrum matches (PSM) of all dependent peptides are shown in a). None of these parameters was affected by TFA incubation in a time-dependent manner. Consistent numbers of ragged-N peptides proof, that no acidic hydrolysis of peptide bonds has occurred. Proportions of distinct dependent peptide modifications relative to the total amount of dependent PSMs were plotted in b), showing that types of peptide modifications remained unaffected as well. Proteins were quantified using the LFQ algorithm in MaxQuant and Pearson correlation coefficients for all sample combinations are shown in c). Comparison of the LFQ intensities for 1 and 60 min TFA incubation prior to sample neutralization is further visualized in a scatter plot (d) with the protein density distribution being color-coded. The TFA incubation time did not alter quantitative reproducibility, as correlation of protein intensities was excellent resulting in Pearson coefficients of at least 0.994.

TFA lysates are prepared for proteolytic digestion by neutralization with a weak base, whose pK_a_ is equal to the optimal pH range of the protease of choice. In such a high capacity buffer system, the desired pH value is stable over a wide range of volume ratios between TFA and base (Fig. S1). TrisBase (pK_a_ =8.1) was chosen to neutralize TFA lysates for tryptic digestion. After neutralization, lysates become slightly turbid as proteins precipitate and form fine particles, but do not tend to aggregate. As neutralization is exothermic, samples instantly heat up to 70 - 80°C. This heat can be exploited to shorten the incubation time needed for reduction and alkylation of disulfide bonds after instant addition of TCEP and CAA to 3 - 5 min.

Proteins in such dispersions are easily accessible for trypsin and can be digested efficiently. However optimal results were obtained after diluting the dispersion 3-10 fold with water to reduce the molarity of salts and the samples' density (data not shown). Tryptic digestion in dispersion was found to proceed similar to in-solution digestion. Optimal enzyme to protein ratio is within 1:20 to 1:100 depending on the protein concentration and 4 -20 h incubation at 37°C are needed to obtain anticipated peptide yields.

The resulting protocol for *Sample Preparation by Easy Extraction and Digestion (SPEED)* consists of three steps, namely acidification, neutralization and digestion. The execution of SPEED is exemplified for *Escherichia coli* and *Staphylococcus aureus* cells as well as for mouse liver tissue in Fig. 2. TFA dissolves *E. coli* and mouse liver tissue completely within 1 and 10 min respectively, while acidification of gram-positive bacteria *S. aureus* forms a cloudy lysate possibly due to insoluble cell wall components (Fig. 2b). Therefore, preparation of gram-positive bacteria requires short microwave irradiation of 10 seconds for efficient digestion. Furthermore TFA does not completely dissolve large collagenous fibers, e.g. from skin preparations, just like detergent-based lysis buffers. Typically this behavior is desirable as abundance of few fiber proteins would otherwise exceed other proteins by far and so impede comprehensive analysis of tissue proteomes. However, if pursued, dissolution of collagen structures can be supported by short microwave irradiation as well.

**Figure 2:**
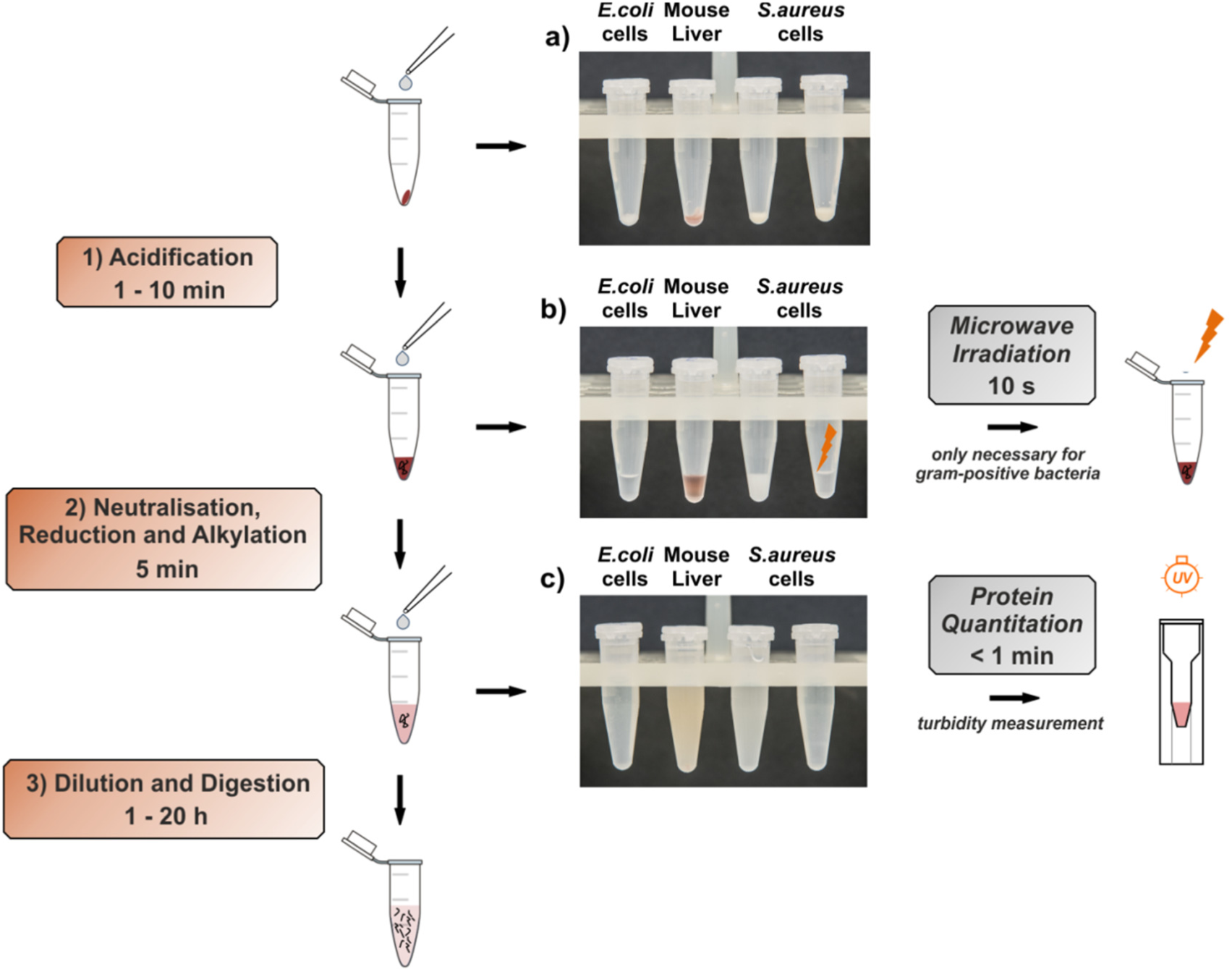
Workflow of Sample Preparation by Easy Extraction and Digestion (SPEED) Sample processing using SPEED consists of three individual steps, namely acidification with pure trifluoroacetic acid (TFA) (1), neutralization with 2M TrisBase (2) and digestion (3). Panels a-c) illustrate steps 1-2 of the procedure for three different sample types. While TFA dissolves *E.coli* cells and mouse liver tissue at room temperature completely in a few minutes, lysis of gram-positive bacteria *S. aureus* requires an additional step of microwave irradiation for 10s (b). After neutralization the samples become slightly turbid (c), as proteins precipitate. Proteins are quantified in the resulting dispersions by turbidity measurement at 360 nm and subsequently digested by addition of trypsin.

After neutralization (Fig. 2c) samples become turbid as proteins precipitate. Proteins are stable in these dispersions during long-term storage at -20°C or -80°C. For protein content determination with conventional methods, proteins need to be solubilized, which is achieved by adding SDS to a sample aliquot prior to performing an assay of choice, e.g. measurements of tryptophan fluorescence ^13^. However, the most straight-forward approach for quantifying proteins in dispersions is by measuring the turbidity, which is an absolute measure for the protein content without the need for any sample manipulation and incubation time. We found that turbidity measurements at 360 nm provided consistent results for different sample types. The data of figure S2 suggest a linear response for protein concentrations between 0.05 – 1 µg/µL (1 AU = 0.79 µg/µL). Protein dispersions are further subjected to digestion after 3-5 fold dilution with water without the need for any purification steps independent of the sample type. Digestion progress can be monitored in real-time using continuous turbidity measurements in a tempered microplate reader as peptides become soluble (Fig. S3).

### Comparison of SPEED with FASP, SP3 and Urea-ISD for the analysis of various sample types

The performance of SPEED was evaluated for the analysis of different sample types in comparison to detergent-based methods (FASP, SP3) ^6,9^ as well as to urea-based in solution digestion ^10^. Sample selection consisted of an easy-to-lyse sample with medium complexity (*E.coli* cells) at different starting amounts (1 and 20 µg protein), an easy-to-lyse sample of high complexity (HeLa cells), a difficult-to-lyse sample of high complexity (mouse lung tissue) and a highly lysis-resistant sample of medium complexity (*B. cereus* cells). All samples were prepared in triplicates and equal peptide amounts of each preparation were analyzed in single-shot LC-MS measurements. The results of SPEED, FASP, SP3 and ISD-Urea were compared by extracting 10 sample preparation dependent parameters from the data, which were categorized into the sections identification, quantification, digestion and modification. Results were normalized relative to the value of the best respective method and visualized in a heatmap to give a general survey of the methods’ performances (Fig. 3).

**Figure 3:**
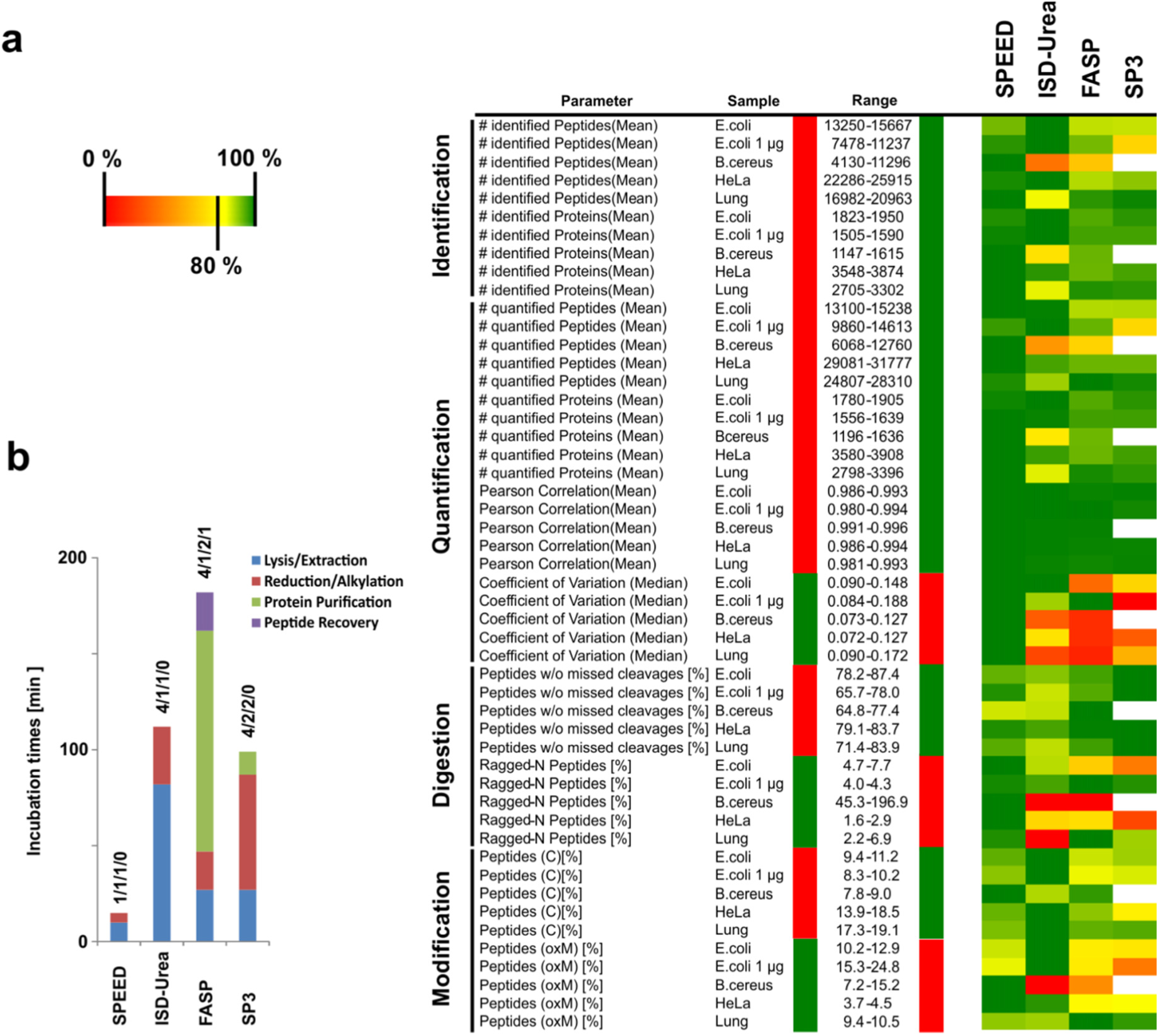
Comparison of SP3, FASP, SPEED and ISD-Urea. Sample preparation by SP3, FASP, SPEED and Urea-based in-solution digestion (ISD-Urea) was compared by triplicate LFQ-based analyses of *E. coli* cells using 20 or 1 µg starting material, *B. cereus* and HeLa cells as well as mouse lung tissue. Results of different parameters sorted according to the sections identification, quantification, digestion and modifications are displayed in a heatmap (a). The best result for each parameter is set to 100 % (dark green) and relative differences between the sample preparation methods are color-coded according to the legend. Red and green surroundings of the range covered by each parameter show if either large or small numbers are regarded as being desirable. As *B. cereus* cells could not be prepared using SP3, the related sections in the heatmap are displayed white. The incubation times and individual numbers of sample handling steps excluding digestion and desalting are compared in (b).

#### Identification

The SPEED protocol enabled identification of most proteins in mouse lung tissue (+3 % compared to FASP), HeLa (+3 % compared to ISD-Urea) and *B. cereus* (+9 % compared to FASP) cells, while ISD-Urea identified most proteins in both analyses of easy-to-lyse *E. coli* cells, exceeding SPEED by 3 % (20 µg starting material) or 1 % (1 µg starting material) respectively. The sample type dependence of the ISD-Urea protocol to effectively extract proteins is illustrated by being the method with the lowest number of protein identifications in both more difficult to lyse sample types, namely mouse lung tissue and *B. cereus*. Concerning protein identifications, FASP and SP3 lagged behind SPEED for all analyzed sample types. Furthermore, we were not able to prepare *B. cereus* for LC-MS analysis using SP3, presumably because cell wall interfered with peptide generation. We observed the same issue for other gram-positive bacteria before, which therefore seems to be an inherent limitation of the original SP3-protocol.

The analysis of low protein quantities usually means that protein extracts are highly diluted, which can impact efficacy of sample processing ^4^. Peptide yields were compared by preparations of *E. coli* cell pellets corresponding to 1 µg protein starting amounts. Cells were lysed in 10 µL of the corresponding buffer and processed by FASP, SP3, SPEED and ISD-Urea. Peptide yield was defined as the peptide intensity relative to its mean value from all measurements injecting 1 µg peptides (20 µg starting material) as calculated by MaxQuant. The median peptide yield was 55 % for SPEED, 44 % for ISD-Urea, 37 % for FASP and 29 % for SP3. Most proteins were identified using ISD-Urea with SPEED (-1 %) lagging slightly and FASP (-5 %) as well as SP3 (-5 %) lagging little further behind.

#### Quantification

Reproducible measurements of protein intensities are essential for quantitative proteome analysis as low variations enhance the statistical power of unambiguous detection of small expression differences. While the protein intensities correlated well for all sample preparation methods with Pearson coefficients exceeding 0.98 for all sample types, coefficients of variation (CV) varied. SPEED samples showed the lowest variations of protein intensities with median CVs of < 9 % for all sample types. ISD-Urea had similar reproducibility compared to SPEED in both analyses of *E. coli* cells, but again suffered from its limited lysis efficacy resulting in enhanced CVs for the analysis of the more difficult-to-lyse sample types, mouse lung tissue (median CV = 16 %) (Fig. S4 a-b) and *B. cereus* (median CV = 12 %). Gene ontology analysis of differentially intense proteins in mouse lung tissue revealed, that membrane proteins are less efficiently prepared for LC-MS by the combination of urea-buffer and ultra-sonication compared to SPEED, FASP and SP3 (Fig. S4 e). As sample preparation using FASP and SP3 resulted in increased CVs compared to SPEED for all sample types and compared to ISD-Urea for all easy-to-lyse sample types, reproducibility of protein quantification seems to benefit from low number of sample handling steps (Fig. 3b). In accordance with this observation, SPEED resulted in the highest number of proteins quantified without missing values for all sample types except *E.coli* (20 µg starting amount), where ISD-Urea exceeded SPEED by 1 %.

#### Digestion

Tryptic digestion of proteins is the fundamental step for bottom-up proteomics sample preparation. The efficacy of proteolysis depends on the source of trypsin, protein-to-enzyme ratio, temperature, and incubation time, which were all kept constant in this study independently of the sample preparation method, as well as on protein concentration and buffer composition, which were variable ^14,15^. The lowest numbers of peptides with missed-cleavage sites were detected in SP3 prepared samples in all experiments (12-22 %). Presumably this finding results from the low digestion volume compared to FASP, SPEED and ISD-Urea. However, higher digestion efficacy is not necessarily associated with lower variability in protein quantitation ^16^. Preparations using SPEED resulted in 2-6 % higher proportions of peptides with missed-cleavage sites, but outperformed SP3 in terms of protein CVs in all experiments. If complete proteolysis would be required, the number of missed-cleavage sites using SPEED can be reduced by increasing the protein concentration or the trypsin-to-protein ratios well as by adding LysC during digestion. The numbers of ragged-N peptides in SPEED preparations were lowest or at least 2^nd^ lowest (+0.07 % relative to total number of peptide IDs) in all preparations. Again, this underlines that peptides are not hydrolyzed by acidification using pure TFA.

#### Modification

Proteins are identified from mass spectra by database searching while usually considering carbamidomethylation of cysteines and oxidation of methionine as possible peptide modifications. FASP, ISD-Urea and SP3 use dithiothreitol (DTT) for reduction and iodoacetamide (IAA) for alkylation of disulfide-bonds consecutively, while SPEED uses tris(2-carboxyethyl)phosphine (TCEP) and chloroacetamide (CAA) simultaneously. The simultaneous use of TCEP and CAA at elevated temperatures reduces sample processing times considerably, however CAA has been found to correlate with increasing numbers of methionine oxidation, although no possible mechanism was proposed ^17^. For method comparison the number of observed modifications was normalized relative to the number of identified peptides in the same sample (Table S2). The largest proportion of cysteine-containing peptides was detected in Urea-ISD prepared samples except for *B. cereus*, however at the expense of increased over-alkylation leading to modification of undesired amino acid residues (Fig. S4). SPEED identified ~ 10 % fewer carbamidomethylated cysteins from HeLa, *E. coli* and mouse lung tissue in comparison to Urea-ISD, but over-alkylation was observed 4-5 fold less frequently. Methionine oxidation was detected at low levels in general (maximum in all samples = 24.6 %) and was further found to be independent from alkylation reagents (Table S2). The differences between various sample types were larger than for different preparations of the same sample and no method was found to lead to increased levels of methionine oxidation in general. This suggests, that unknown factors influence oxidation. Therefore, at least when reduction and alkylation reagents are solubilized immediately before use, utilization of TCEP and CAA had no unwanted side-effects compared to DTT and IAA.

Other frequently observed sample preparation related modifications in proteomics are deamidation, carbamylation, loss of ammonia and oxidation of residues excluding methionine ^18^. Results of a dependent peptide search revealed, that the lowest total numbers of these unintentionally introduced modifications were observed for SP3 and the highest total numbers for ISD-Urea in all sample types (Fig. S5). SPEED and FASP were on pair while introducing 1-2 % more artificial modifications relative to the total number of identified peptides compared to SP3. An open search analysis of mouse lung tissue data using MSFragger^19^ further revealed that SPEED does not add any type of unannotated modification to peptides (data not shown).

## DISCUSSION

Comprehensive, accurate and deep proteome analysis relies on the reproducible generation of peptides from the entity of proteins of a given sample type. SPEED offers some inherent benefits over common methods for this purpose. It is a minimal strategy for universal proteomics sample preparation based on a fundamentally different chemistry compared to existing methods. SPEED combines the straight-forward nature of in-solution digestion with a highly efficient one-step extraction even of membrane proteins from tissue samples or lysis-resistant gram-positive bacteria. TFA extraction circumvents the use of detergents and chaotropic agents as well as of physical disruption methods for lysis and protein extraction. This enables sample processing in a truly one pot manner including protein extraction by simple acidification. The SPEED sample preparation protocol generates clear peptide solutions even from tissue samples without removal of any sample material and therefore without protein losses. Analysis of *E. coli* lysates with prolonged TFA-incubation times of up to 1 h at room temperature revealed, that TFA does not hydrolyze or modify protein primary structures. Acidification with TFA is further known to reliably inactivate pathogenic microorganisms including bacterial endospores from *Bacillus anthracis* and is therefore well suited for the sample processing of infectious samples, including highly pathogenic agents ^20^. The rapid processing, high performance and broad applicability of the SPEED protocol is realized solely on the smart use of chemicals and requires no special equipment or even centrifuges. It is an inexpensive method with costs of ~ 1 € per sample including pipette tips and microcentrifuge tubes. Furthermore, the protocol can easily be accomplished by non-experts and reduces hands-on processing times down to 15 min including lysis and protein quantitation. Protein content of SPEED lysates is determined by turbidity measurement, which does not require any sample manipulation and therefore enables complete recovery of proteins. Furthermore turbidity measurements allow real-time monitoring of the digestion progress, which could be used as a quality criterion prior to LC-MS measurements. Due to its inherent benefits, SPEED has the potential to be widely adopted in the proteomics community as a rapid, low-cost, detergent-free and universal sample preparation method. In the future, SPEED processing can be transferred to liquid-handling stations starting from lysis of even tissue by simply pipetting up and down and ending by on-line monitoring of the digestion process using turbidity measurements.

SPEED is not only universally applicable but also turned out to be superior compared to the widely adopted methods FASP, SP3 and ISD-Urea for quantitative proteome analysis of various sample types. SPEED outperformed FASP and SP3 considering the number of identified proteins for all samples analyzed and ISD-Urea for HeLa, *B. cereus* and mouse lung tissue, respectively. Furthermore SPEED and ISD-Urea were found to provide increased peptide yields from 1 µg preparations of a diluted *E. coli* lysate (0.1 µg/µL) resulting in elevated numbers of protein identifications. We conclude that the increased peptide yield of SPEED and ISD-Urea resulted from the fewer handling steps compared to FASP and SP3, which require removal of detergents prior to digestion and are therefore prone to protein losses at low starting amounts. Reproducibility of protein quantification benefitted from low number of sample handling steps as well since sample preparation using FASP and SP3 resulted in increased CVs compared to SPEED for all sample types and compared to ISD-Urea for all easy-to-lyse sample types. Sample preparation using SPEED resulted not only in the best quantitative reproducibility, but also in the highest number of proteins quantified without missing values for all sample types except *E. coli* (20 µg starting amount), where ISD-Urea exceeded SPEED by 1 %. However, the high performance of ISD-Urea for easy-to-lyse *E. coli* cells was compensated by the results of the analysis of mouse lung tissue and *B. cereus*. Membrane proteins were less efficiently extracted in lung tissue by the combination of urea-buffer and ultra-sonication compared to the detergent-based methods FASP and SP3 as well as to SPEED. ISD-Urea treatment was further not suitable for effective lysis of highly-resistant *B. cereus* cells, which decreased the number of protein identifications by 30 % and of peptide identifications by 64 % compared to SPEED.

The comparative analysis of detergent-based (FASP, SP3), chaotropic agent-based (ISD-Urea) and acid-based (SPEED) sample preparation protocols for the quantitative proteome analysis of different sample types clearly demonstrates the benefits of low number of sample handling steps for improving quantitative reproducibility, number of quantified proteins and peptide yields from low starting amounts. SPEED combines the straight-forward nature of in-solution digestion with highly efficient one-step extraction by acidification. This protocol thus requires the lowest number of sample handling steps and the shortest hands-on time. SPEED was found to be the best suited protocol tested for quantifying the largest number of proteins with highest accuracy for a wide selection of sample types and can therefore be regarded as a universal sample preparation method for bottom-up proteomics.

## EXPERIMENTAL PROCEDURES

### Sample Materials

Tryptic Soy Agar (TSA) ReadyPlates^TM^ (Merck, Darmstadt, Germany) were inoculated with *E. coli K-12* (DSM 3871), *S. aureus* (DSM 4910) or *B. cereus* (ATCC^®^ 10987) and incubated at 37°C overnight. Cells were harvested using an inoculating loop and washed in 2 x 1 mL phosphate-buffered saline (PBS) for 5 min at 4,000 x g and 4°C. Cells were aliquoted, again pelleted and kept at -80°C until lysis.

HeLa cells (ATCC^®^ CCL-2™) were cultivated in DMEM supplemented with 10 % FCS and 2 mM L–Glutamine at 37°C and harvested at 90 % confluency by scraping. Cells were washed in 2 x 2 mL phosphate-buffered saline (PBS) for 8 min at 400 x g and 4°C, aliquoted, again pelleted and kept at -80°C until lysis.

BALB/c mouse lung and liver were obtained from the central laboratory animal facility (MF3, Robert Koch-Institute, Berlin, Germany), washed 3 x by dipping into 10 mL ice-cold PBS, cut into slices, aliquoted, flash frozen with liquid nitrogen and kept at -80°C until lysis. After addition of lysis buffer (FASP, SP3, Urea-ISD) mouse lung samples were transferred to Lysing Matrix D Tubes (MP Biomedicals, Santa Ana, California, USA) and subjected to grinding.

### Sample Preparation by Easy Extraction and Digestion (SPEED)

Samples were resuspended in trifluoroacetic acid (TFA) (Uvasol^®^ for spectroscopy, Merck, Darmstadt, Germany) (sample/TFA 1:4 (v/v) or 10 µL for the E.coli 1 µg experiment) and incubated at room temperature for 2 (*E.coli, B. cereus*, HeLa) or 10 (mouse lung tissue) min. *B. cereus* samples were further irradiated for 10 s at 800 W using a microwave oven. Samples were neutralized with 2M TrisBase using 10 x volume of TFA and further incubated at 95°C for 5 min after adding Tris(2-carboxyethyl)phosphine (TCEP) to a final concentration of 10 mM and 2-Chloroacetamide (CAA) to a final concentration of 40 mM. Protein concentrations were determined by turbidity measurements at 360 nm (1 AU = 0.79 µg/µL) using GENESYS™ 10S UV-Vis Spectrophotometer (Thermo Fisher Scientific, Waltham, Massachusetts, USA) and adjusted to 0.25 µg/µL using a 10:1 (v/v) mixture of 2M TrisBase and TFA and then diluted 1:5 with water. Digestion was carried out for 20 h at 37°C using Trypsin Gold, Mass Spectrometry Grade (Promega, Fitchburg, WI, USA) at a protein/enzyme ratio of 50:1.

### Filter-aided Sample Preparation (FASP)

Samples were suspended in 4 % SDS, 100 mM Tris/HCl, 100 mM DTT, pH 7.6 (sample/buffer 1:10 (v/v) or 10 µL for the *E.coli* 1 µg experiment), incubated at 95°C for 5 min and further sonicated for 10 (*E.coli*, HeLa) or 15 (*B. cereus*, mouse lung tissue) cycles a 30 s at high intensity level and 4°C using Bioruptor^®^Plus (Diagenode, Liege, Belgium). Samples were clarified by centrifugation at 16,000 x g for 5 min und processed using Microcon-30kDa Centrifugal Filter Units (Merck, Darmstadt, Germany) according to the Filter-aided Sample Preparation (FASP) protocol of Wisniewski et al. ^6^ with proteins being digested for 20 h at 37°C using Trypsin Gold, Mass Spectrometry Grade (Promega, Fitchburg, WI, USA) at a protein/enzyme ratio of 50:1.

### Single-Pot Solid-Phase-enhanced Sample Preparation (SP3)

Samples were suspended in 1 % SDS, 1X cOmplete Protease Inhibitor Cocktail (Roche, Basel, Switzerland), 50 mM HEPES buffer, pH 8.5 (sample/buffer 1:10 (v/v) or 10 µL for the *E.coli* 1 µg experiment), incubated at 95°C for 5 min and further sonicated for 10 (*E.coli*, HeLa) or 15 (*B. cereus*, mouse lung tissue) cycles a 30 s at high intensity level and 4°C using Bioruptor^®^Plus (Diagenode, Liege, Belgium). Samples were further processed according to the Single-Pot Solid-Phase-enhanced Sample Preparation (SP3) method of Hughes et al. ^9^, except that proteins were bound to the paramagnetic beads at 70 % ACN without acidification as prescribed by Sielaff et al. ^4^. Digestion was carried out for 20 h at 37°C using Trypsin Gold, Mass Spectrometry Grade (Promega, Fitchburg, WI, USA) at a protein/enzyme ratio of 50:1.

### Urea-based in-solution digestion (Urea-ISD)

Samples were suspended in 8M urea, 50mM Tris-HCl, 5mM DTT, pH 8 (sample/buffer 1:10 (v/v) or 10 µL for the *E.coli* 1 µg experiment) and sonicated for 10 (*E.coli*, HeLa) or 15 (*B. cereus*, mouse lung tissue) cycles a 30 s at high intensity level and 4°C using Bioruptor^®^Plus (Diagenode, Liege, Belgium). Samples were further incubated for 1 h at 37°C and clarified by centrifugation at 16,000 x g for 5 min. IAA was added to a final concentration of 15 mM and samples were alkylated for 30 min at room temperature in the dark. Urea was diluted with 50mM Tris-HCl (pH 8) to 1 M, Trypsin Gold, Mass Spectrometry Grade (Promega, Fitchburg, WI, USA) was added at a protein/enzyme ratio of 50:1 and proteins were digested for 20 h at 37°C.

### Measurements of Tryptophan Fluorescence

Protein concentrations were determined by measuring the tryptophan fluorescence at an emission wavelength of 350 nm using 295 nm for excitation with an Infinite^®^ M1000 PRO microplate reader (Tecan, Maennedorf, Switzerland) ^13^. The tryptophan content of each sample was determined using a standard curve ranging from 0.1 – 0.9 µg tryptophan and assuming a tryptophan weight content of 1.3 % in the samples. 20 µg protein of each sample type was further digested except for the *E.coli* experiment with low starting amount (*E.coli* 1µg). Therefore, protein content of a cell suspension was determined after TFA-based lysis and cells from the suspension volume equivalent to 1 µg protein were aliquoted and pelleted.

### Turbidity Measurements

Turbidity of SPEED-lysates was measured at 360 nm using a GENESYS™ 10S UV-Vis Spectrophotometer (Thermo Fisher Scientific, Waltham, Ma, USA). If the initial protein concentration was above 1 µg/µL, samples were diluted with a 10:1 (v/v) mixture of 2M TrisBase and TFA. Protein concentrations were calculated using the correlation of 1 AU = 0.79 µg/µL. For experiments analyzing the digestion progress by real-time monitoring, an Infinite^®^ M1000 PRO microplate reader (Tecan, Maennedorf, Switzerland) was used. The microplate reader was tempered to 37°C and turbidity was measured at 360 nm every 5 min.

### Peptide Desalting

Peptides generated using FASP, SPEED and Urea-ISD were desalted using 200 µL StageTips packed with three Empore™ SPE Disks C18 (3M Purification, Inc., Lexington, USA) according to Rappsilber et al. ^21^ and concentrated using a vacuum concentrator. Samples were resuspended in 20 µL 0.1 % formic acid and peptides were quantified by measuring the absorbance at 280 nm using a Nanodrop 1000 (Thermo Fisher Scientific, Rockford, IL, USA).

### Liquid chromatography and mass spectrometry

Peptides were analyzed on an EASY-nanoLC 1200 (Thermo Fisher Scientific, Bremen, Germany) coupled online to a Q Exactive™ Plus mass spectrometer (Thermo Fisher Scientific, Bremen, Germany). 1 µg peptides were separated on a 50 cm Acclaim™ PepMap™ column (75 μm i.d., 100 Å C18, 2 μm; Thermo Fisher Scientific, Bremen, Germany) using a linear 120 (*E.coli, B. cereus*) or 180 (HeLa, mouse lung tissue) min gradient of 3 to 28 % acetonitrile in 0.1 % formic acid at 200 nL/min flow rate. Column temperature was kept at 40°C using a butterfly heater (Phoenix S&T, Chester, PA, USA). The Q Exactive™ Plus was operated in a data-dependent manner in the m/z range of 300 – 1,650. Full scan spectra were recorded with a resolution of 70,000 using an automatic gain control (AGC) target value of 3 × 10^6^ with a maximum injection time of 20 ms. Up to the 10 most intense 2^+^ - 5^+^ charged ions were selected for higher-energy c-trap dissociation (HCD) with a normalized collision energy (NCE) of 25 %. Fragment spectra were recorded at an isolation width of 2 Th and a resolution of 17,500@200m/z using an AGC target value of 1 × 10^5^ with a maximum injection time of 50 ms. The minimum MS² target value was set to 1 × 10^4^. Once fragmented, peaks were dynamically excluded from precursor selection for 30 s within a 10 ppm window. Peptides were ionized using electrospray with a stainless steel emitter, I.D. 30 µm, (Proxeon, Odense, Denmark) at a spray voltage of 2.0 kV and a heated capillary temperature of 275°C.

## Data analysis

Mass spectra were analyzed using MaxQuant (Version 1.5.1.2) ^22^. At first, parent ion masses were recalibrated using the 'software lock mass’ option before the MS² spectra were searched using the Andromeda algorithm against sequences from the complete proteomes of either *homo sapiens* (UP000005640), *E.coli K-12* (UP000000625), *B.cereus strain ATCC 10987* (UP000002527) or *mus musculus* (UP000000589), which were downloaded from UniProt. Spectra were searched with a tolerance of 4.5 ppm in MS^1^ and 20 ppm in HCD MS² mode, strict trypsin specificity (KR not P) and allowing up to two missed cleavage sites. Cysteine carbamidomethylation was set as a fixed modification and methionine oxidation as well as N-terminal acetylation of proteins as variable modifications. The false discovery rate (FDR) was set to 1 % for peptide and protein identifications. Dependent peptides with previously unconsidered modifications were identified using 1 % FDR as well. Identifications were transferred between samples using the 'match between run’ option within a match window of 0.7 min and an alignment window of 20 min. Protein intensities for label-free quantification (LFQ) were calculated separately for each sample preparation method. The analysis was once repeated for each experiment allowing semi-specific tryptic digestion at the peptide N-termini.

Analysis of the MaxQuant results was done in Persues (Version 1.5.0.31) ^23^. At first, reverse hits, contaminants and proteins only identified by site were removed. Afterwards protein related parameters (identified proteins, quantified proteins, Pearson correlation coefficients, coefficients of variation), peptide related parameters (identified peptides, quantified peptides, missed cleavage sites, ragged-N peptides, modified peptides (C, oxM)) and dependent peptide related parameters (distribution of sample preparation-related peptide modifications) were extracted from the respective .txt files. Only peptides and proteins without missing values were considered for quantification.

---

**Fig. S1:**
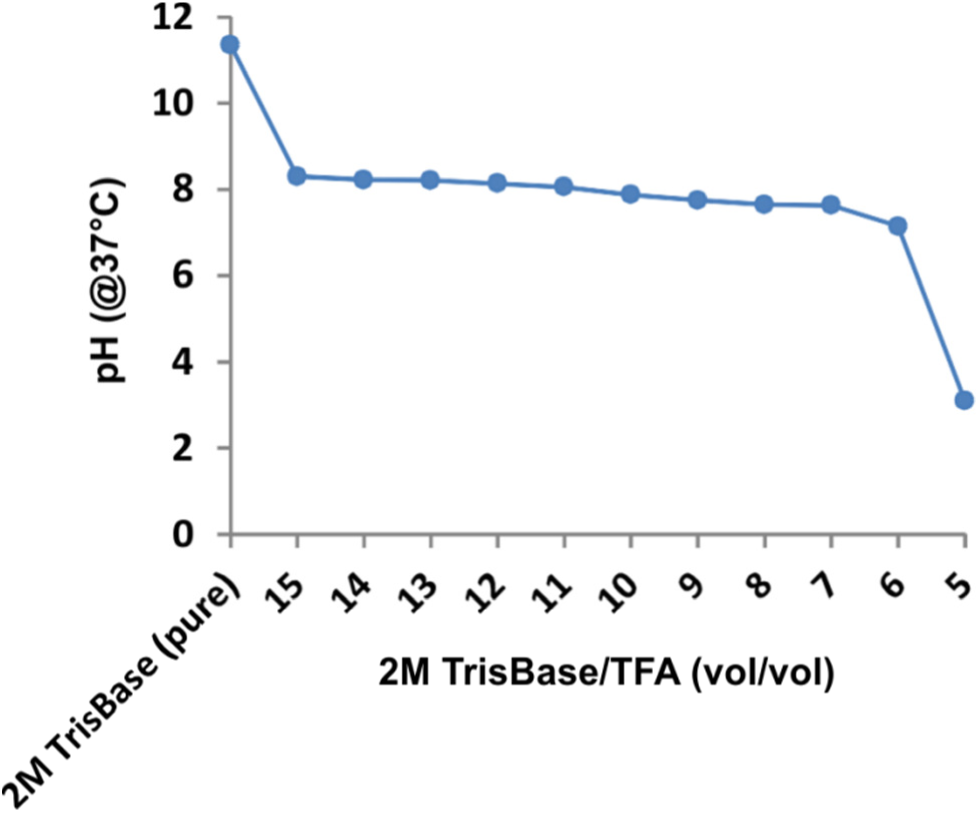
pH Stability of the SPEED buffer system. TrisBase was chosen to neutralize TFA for SPEED-based sample processing. The pH value of TFA/TrisBase mixtures is stable at the pK_a_ = 8.1 of TrisBase over a wide range of volume ratios and is therefore well suited to enable tryptic digestion of proteins.

**Fig. S2:**
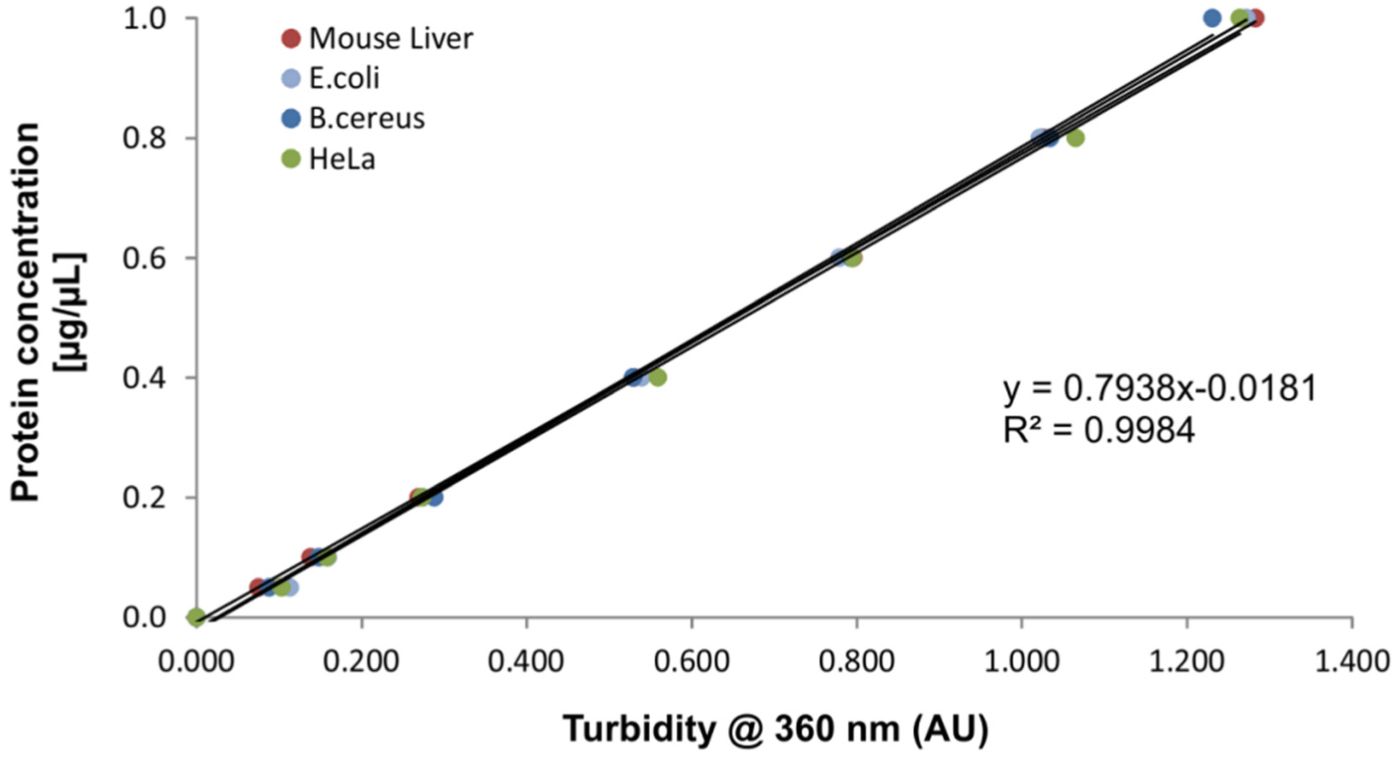
Determination of protein concentrations using turbidity measurements. Turbidity of various SPEED-prepared sample types was measured at 360 nm in the concentration range of 0.05 – 1 µg/µL. Tryptophan flourescence measurement was used as the reference method for protein content determination. Turbidity was independent of the sample type and correlates linearily with protein concentration (1 AU = 0.79 µg/µL).

**Fig. S3:**
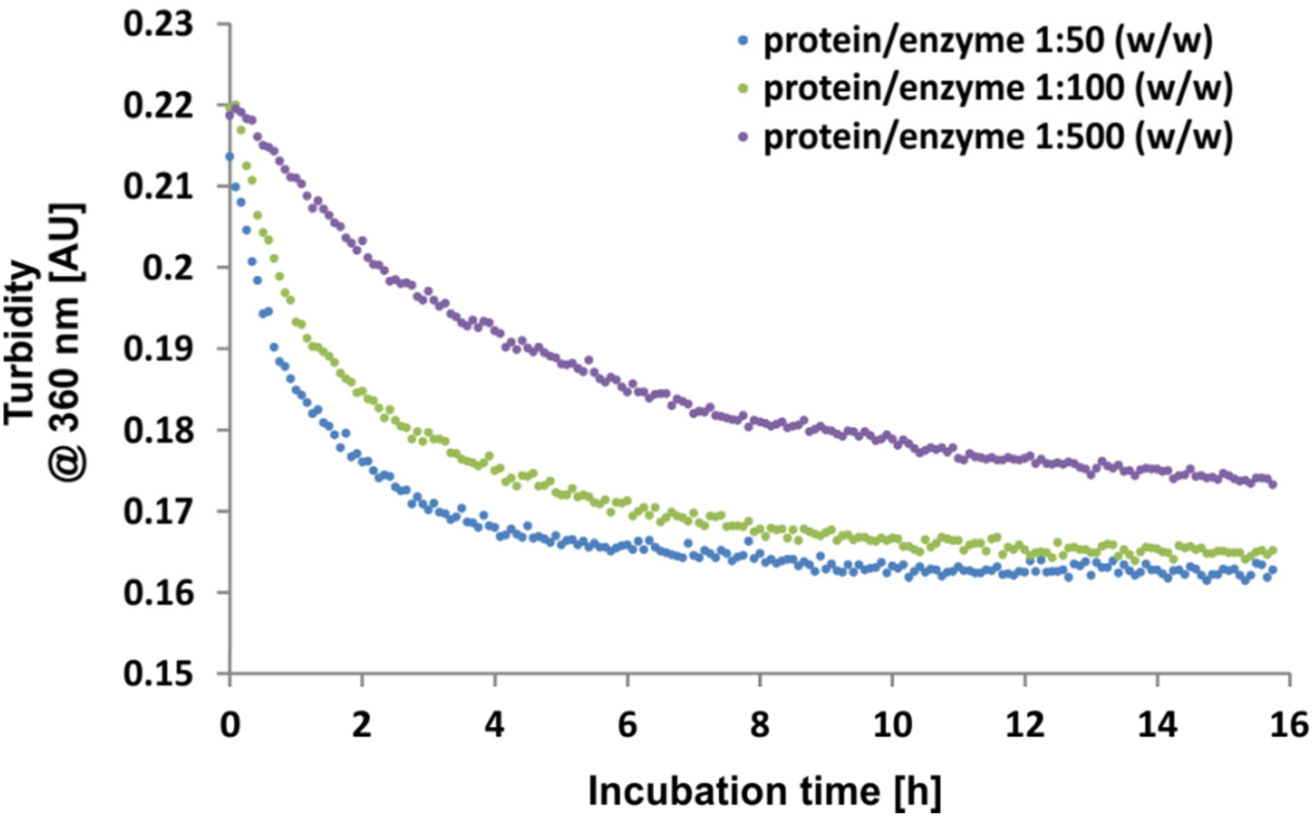
Real-time monitoring of tryptic digestion using turbidity measurements. SPEED-prepared E.coli lysates (20 µg) were digested at different protein/enzyme (w/w) ratios. Turbidity was measured in real-time at 360 nm using a microplate reader tempered to 37°C. Progress of tryptic protein digestion correlates with decreasing turbidity as generated peptides become water-soluble.

**Fig. S4:**
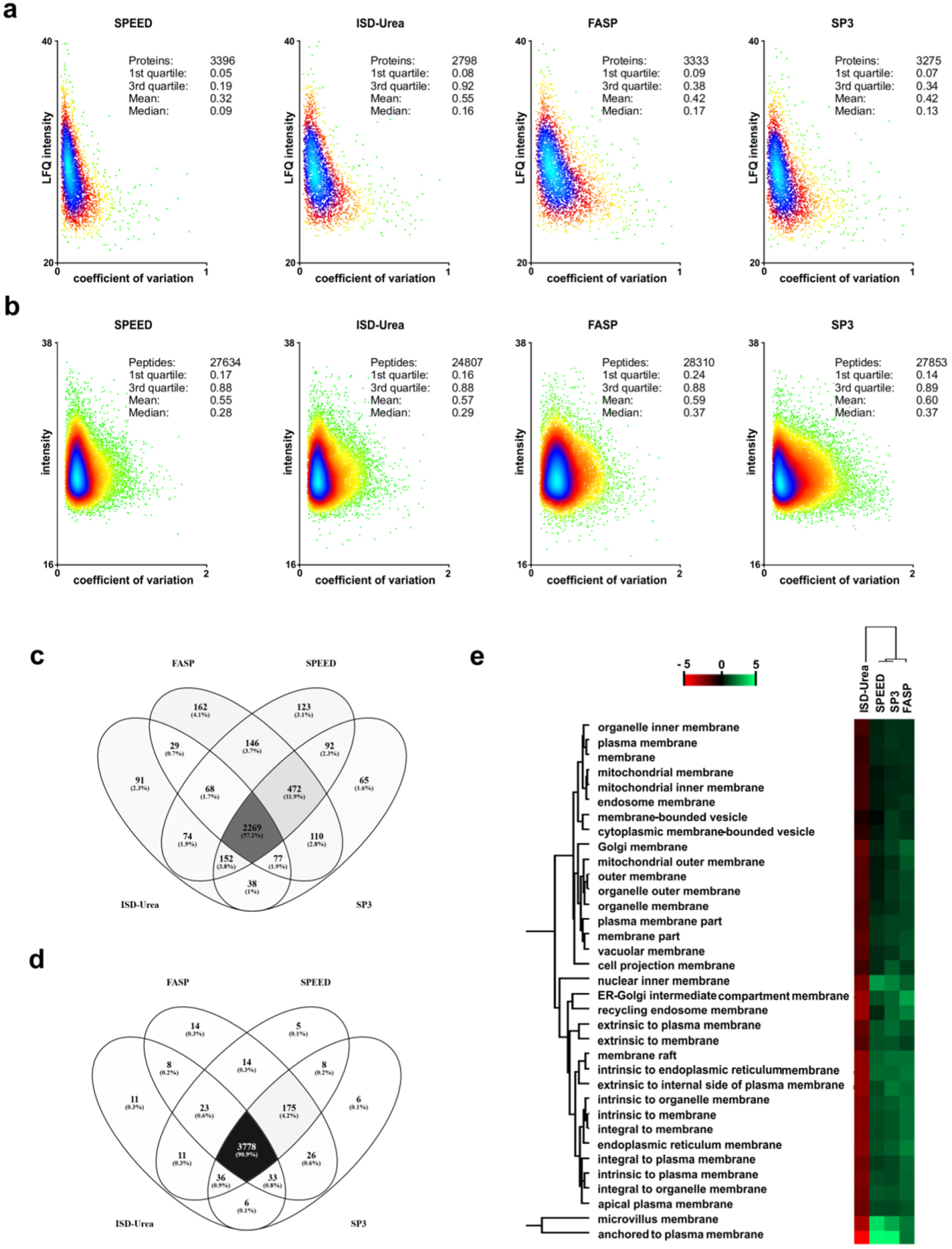
Comparison of SP3, FASP, SPEED and ISD-Urea for the analysis of mouse lung tissue. Mouse lung tissue was prepared in triplicates using SP3, FASP, SPEED and ISD-Urea. Proteins were quantified from the LC-MS/MS data using LFQ-algorithm in MaxQuant. Coefficients of variation (CV) are plotted against their corresponding intensities for proteins (a) and peptides (b) with the density distributions being color-coded. Protein quantifications (c) and identifications (d) of the different sample preparation methods are compared in Venn diagrams. Mean Tukey’s honestly significant difference (THSD) of proteins with cellular component ontologies associated with membranes are compared in the heatmap (e)

**Fig. S5:**
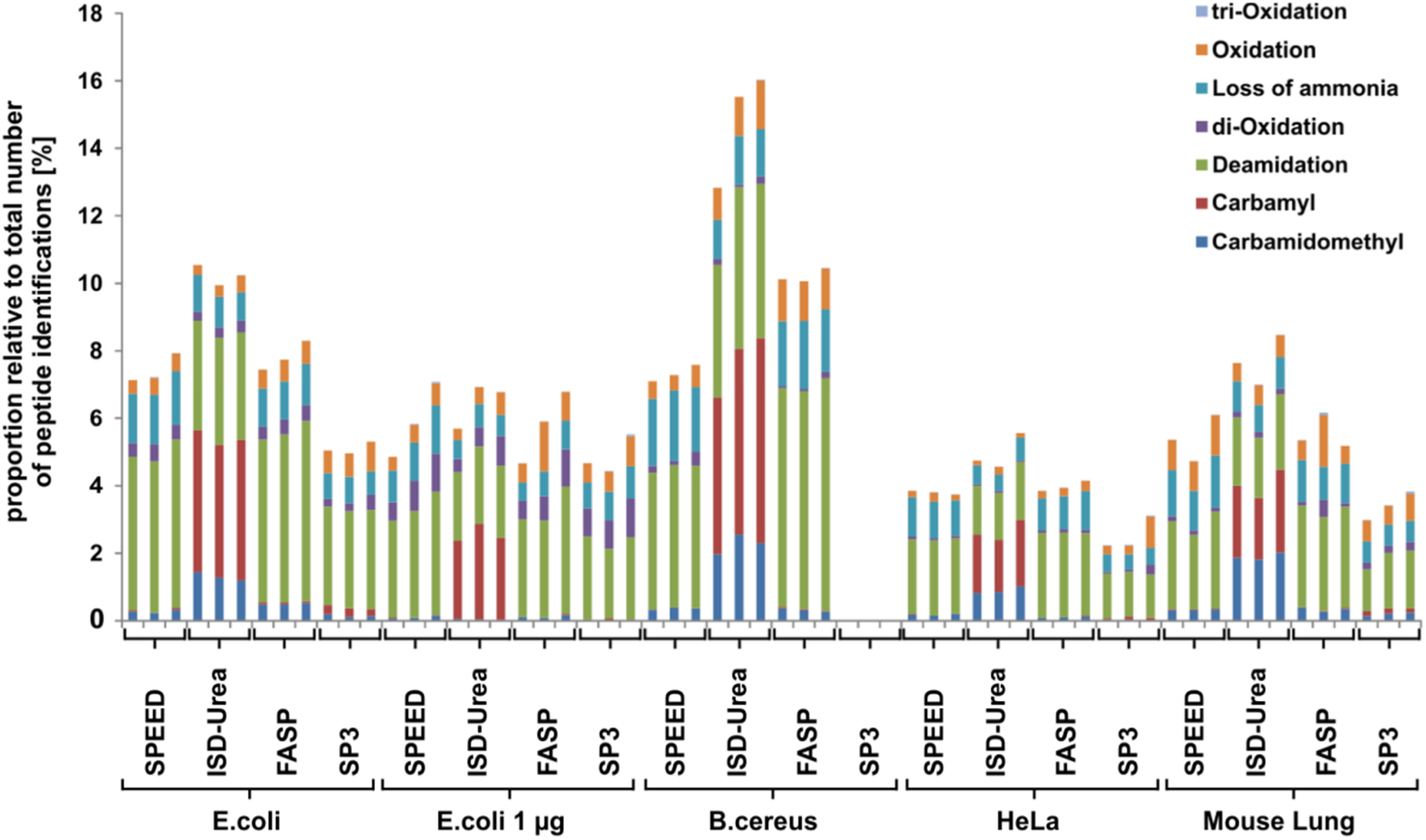
Comparison of dependent peptide IDs between SP3, FASP, SPEED and ISD-Urea. Dependent peptide identifications of modifications known to be introduced during sample preparation are displayed relative to the total number of peptide identiciations for each analyzed sample. The proportion of individual modification types are color-coded.

**Table:S1:** Absolute numerical values of SP3, FASP, SPEED and ISD-Urea comparison.

|  | Parameter | Sample | SPEED | ISD-Urea | FASP | SP3 |
| --- | --- | --- | --- | --- | --- | --- |
| Identification | # identified Peptides (Mean) | E.coli | 14219 | 15667 | 13311 | 13250 |
|  | # identified Proteins (Mean) | E.coli | 1899 | 1950 | 1823 | 1884 |
|  | # quantified Peptides (Mean) | E.coli | 15238 | 15184 | 13100 | 13141 |
|  | # quantified Proteins (Mean) | E.coli | 1883 | 1905 | 1780 | 1844 |
| Quantification | Pearson Correlation (Mean) | E.coli | 0.99 | 0.99 | 0.99 | 0.99 |
|  | Coefficient of Variation (Median) | E.coli | 0.09 | 0.09 | 0.15 | 0.12 |
|  | Peptides w/o missed cleavages (Mean) | E.coli | 80.64 | 78.22 | 82.73 | 87.41 |
|  | Ragged-N Peptides (Mean) | E.coli | 675 | 851 | 857 | 1015 |
|  | Ragged-N Peptides [%] | E.coli | 4.75 | 5.43 | 6.44 | 7.66 |
|  | Peptides (C) (Mean) | E.coli | 1473 | 1755 | 1257 | 1299 |
|  | Peptides (C) [%] | E.coli | 10.36 | 11.20 | 9.44 | 9.81 |
|  | Peptides (oxM) (Mean) | E.coli | 11.76 | 10.15 | 12.75 | 12.94 |
|  | Chemical Modifications (Mean) | E.coli | 1055 | 1605 | 1040 | 675 |
| Digestion | Chemical Modifications [%] | E.coli | 7.42 | 10.24 | 7.81 | 5.09 |
|  | Parameter | Sample | SPEED | ISD-Urea | FASP | SP3 |
| Identification | # identified Peptides (Mean) | E.coli 1µg | 10871 | 11237 | 10207 | 7478 |
|  | # identified Proteins (Mean) | E.coli 1µg | 1574 | 1590 | 1505 | 1506 |
|  | # quantified Peptides (Mean) | E.coli 1µg | 13963 | 14613 | 13440 | 9860 |
|  | # quantified Proteins (Mean) | E.coli 1µg | 1639 | 1628 | 1556 | 1557 |
| Quantification | Pearson Correlation (Mean) | E.coli 1µg | 0.99 | 0.99 | 0.99 | 0.98 |
|  | Coefficient of Variation (Median) | E.coli 1µg | 0.08 | 0.09 | 0.08 | 0.19 |
|  | Peptides w/o missed cleavages (Mean) | E.coli 1µg | 75.93 | 65.69 | 72.81 | 78.00 |
|  | Ragged-N Peptides (Mean) | E.coli 1µg | 446 | 478 | 411 | 310 |
|  | Ragged-N Peptides [%] | E.coli 1µg | 4.10 | 4.25 | 4.03 | 4.15 |
|  | Peptides (C) (Mean) | E.coli 1µg | 988 | 1147 | 848 | 635 |
|  | Peptides (C) [%] | E.coli 1µg | 9.09 | 10.21 | 8.31 | 8.49 |
|  | Peptides (oxM) (Mean) | E.coli 1µg | 18.09 | 15.30 | 19.33 | 24.84 |
|  | Chemical Modifications (Mean) | E.coli 1µg | 649 | 726 | 585 | 366 |
| Digestion | Chemical Modifications [%] | E.coli 1µg | 5.97 | 6.46 | 5.73 | 4.89 |
|  | Parameter | Sample | SPEED | ISD-Urea | FASP | SP3 |
| Identification | # identified Peptides (Mean) | B.cereus | 11296 | 4130 | 6996 | n.d. |
|  | # identified Proteins (Mean) | B.cereus | 1615 | 1147 | 1480 | n.d. |
|  | # quantified Peptides (Mean) | B.cereus | 12760 | 6068 | 8463 | n.d. |
|  | # quantified Proteins (Mean) | B.cereus | 1636 | 1196 | 1494 | n.d. |
| Quantification | Pearson Correlation (Mean) | B.cereus | 1.00 | 0.99 | 0.99 | n.d. |
|  | Coefficient of Variation (Median) | B.cereus | 0.07 | 0.12 | 0.13 | n.d. |
|  | Peptides w/o missed cleavages (Mean) | B.cereus | 64.81 | 65.71 | 77.44 | n.d. |
|  | Ragged-N Peptides (Mean) | B.cereus | 5122 | 8132 | 6857 | n.d. |
|  | Ragged-N Peptides [%] | B.cereus | 45.35 | 196.91 | 98.00 | n.d. |
|  | Peptides (C) (Mean) | B.cereus | 1018 | 321 | 605 | n.d. |
|  | Peptides (C) [%] | B.cereus | 9.01 | 7.78 | 8.65 | n.d. |
|  | Peptides (oxM) (Mean) | B.cereus | 7.24 | 15.22 | 11.28 | n.d. |
|  | Chemical Modifications (Mean) | B.cereus | 826 | 608 | 715 | n.d. |
| Digestion | Chemical Modifications [%] | B.cereus | 7.32 | 14.72 | 10.21 | n.d. |
|  | Parameter | Sample | SPEED | ISD-Urea | FASP | SP3 |
| Identification | # identified Peptides (Mean) | HeLa | 25399 | 25915 | 22286 | 23036 |
|  | # identified Proteins (Mean) | HeLa | 3874 | 3744 | 3548 | 3660 |
|  | # quantified Peptides (Mean) | HeLa | 31777 | 29979 | 29081 | 29106 |
|  | # quantified Proteins (Mean) | HeLa | 3908 | 3727 | 3580 | 3706 |
| Quantification | Pearson Correlation (Mean) | HeLa | 0.99 | 0.99 | 0.99 | 0.99 |
|  | Coefficient of Variation (Median) | HeLa | 0.07 | 0.09 | 0.13 | 0.12 |
|  | Peptides w/o missed cleavages (Mean) | HeLa | 81.65 | 79.15 | 83.59 | 83.73 |
|  | Ragged-N Peptides (Mean) | HeLa | 413 | 559 | 473 | 665 |
|  | Ragged-N Peptides [%] | HeLa | 1.62 | 2.16 | 2.12 | 2.89 |
|  | Peptides (C) (Mean) | HeLa | 4319 | 4790 | 3661 | 3197 |
|  | Peptides (C) [%] | HeLa | 17.01 | 18.48 | 16.43 | 13.88 |
|  | Peptides (oxM) (Mean) | HeLa | 3.71 | 3.83 | 4.52 | 4.45 |
|  | Chemical Modifications (Mean) | HeLa | 966 | 1281 | 888 | 580 |
| Digestion | Chemical Modifications [%] | HeLa | 3.80 | 4.94 | 3.99 | 2.52 |
|  | Parameter | Sample | SPEED | ISD-Urea | FASP | SP3 |
| Identification | # identified Peptides (Mean) | Lung | 20963 | 16982 | 20240 | 20750 |
|  | # identified Proteins (Mean) | Lung | 3302 | 2705 | 3210 | 3189 |
|  | # quantified Peptides (Mean) | Lung | 27634 | 24807 | 28310 | 27853 |
|  | # quantified Proteins (Mean) | Lung | 3396 | 2798 | 3333 | 3275 |
| Quantification | Pearson Correlation (Mean) | Lung | 0.99 | 0.99 | 0.98 | 0.99 |
|  | Coefficient of Variation (Median) | Lung | 0.09 | 0.16 | 0.17 | 0.13 |
|  | Peptides w/o missed cleavages (Mean) | Lung | 78.00 | 71.45 | 79.70 | 83.90 |
|  | Ragged-N Peptides (Mean) | Lung | 466 | 1179 | 439 | 507 |
|  | Ragged-N Peptides [%] | Lung | 2.22 | 6.94 | 2.17 | 2.44 |
|  | Peptides (C) (Mean) | Lung | 3622 | 3251 | 3580 | 3708 |
|  | Peptides (C) [%] | Lung | 17.28 | 19.14 | 17.69 | 17.87 |
|  | Peptides (oxM) (Mean) | Lung | 10.43 | 10.53 | 9.36 | 9.74 |
|  | Chemical Modifications (Mean) | Lung | 1127 | 1306 | 1128 | 706 |
| Digestion | Chemical Modifications [%] | Lung | 5.37 | 7.69 | 5.57 | 3.40 |

**Table S2:** Comparison of reduction and alkylation efficacy of disulfide bonds between SP3, FASP, SPEED and ISD-Urea.

| Parameter | Sample | ISD-Urea | SPEED | FASP | SP3 |
| --- | --- | --- | --- | --- | --- |
| *Peptides (C) [%] (Mean)* | HeLa | 18.5 (+/- 0.5) | 17.0 (+/- 0.1) | 16.4 (+/- 0.5) | 13.9 (+/- 0.5) |
| *Peptides (oxM) [%] (Mean)* | HeLa | 3.8 (+/- 0.3) | 3.7 (+/- 0.0) | 4.5 (+/- 0.3) | 4.4 (+/- 0.2) |
| *Carbamidomethyl (excl. C)* | HeLa | 233 (+/- 12) | 45 (+/- 6) | 23 (+/- 3) | 7 (+/- 2) |
| Parameter | Sample | ISD-Urea | SPEED | FASP | SP3 |
| *Peptides (C) [%] (Mean)* | E.coli | 11.2 (+/- 0.1) | 10.4 (+/- 0.1) | 9.4 (+/- 0.1) | 9.8 (+/- 0.4) |
| *Peptides (oxM) [%] (Mean)* | E.coli | 10.2 (+/- 0.5) | 11.8 (+/- 0.4) | 12.7 (+/- 0.5) | 12.9 (+/- 1.6) |
| *Carbamidomethyl (excl. C)* | E.coli | 205 (+/- 28) | 39 (+/- 7) | 66 (+/- 1) | 20 (+/- 7) |
| Parameter | Sample | ISD-Urea | SPEED | FASP | SP3 |
| *Peptides (C) [%] (Mean)* | E.coli 1µg | 10.2 (+/- 0.2) | 9.1 (+/- 0.4) | 8.3 (+/- 0.3) | 8.5 (+/- 0.5) |
| *Peptides (oxM) [%] (Mean)* | E.coli 1µg | 15.3 (+/- 1.1) | 18.1 (+/- 1.0) | 19.3 (+/- 2.4) | 24.8 (+/- 4.0) |
| *Carbamidomethyl (excl. C)* | E.coli 1µg | 4 (+/- 2) | 11 (+/- 5) | 12 (+/- 4) | 1 (+/- 1) |
| Parameter | Sample | ISD-Urea | SPEED | FASP | SP3 |
| *Peptides (C) [%] (Mean)* | B.cereus | 7.8 (+/- 0.6) | 9.0 (+/- 0.0) | 8.6 (+/- 0.1) | n.d. |
| *Peptides (oxM) [%] (Mean)* | B.cereus | 15.2 (+/- 2.3) | 7.2 (+/- 0.3) | 11.3 (+/- 0.3) | n.d. |
| *Carbamidomethyl (excl. C)* | B.cereus | 94 (+/- 11) | 41 (+/- 4) | 22 (+/- 4) | n.d. |
| Parameter | Sample | ISD-Urea | SPEED | FASP | SP3 |
| *Peptides (C) [%] (Mean)* | Mouse Lung | 19.1 (+/- 0.4) | 17.3 (+/- 0.3) | 17.7 (+/- 0.4) | 17.9 (+/- 0.2) |
| *Peptides (oxM) [%] (Mean)* | Mouse Lung | 10.5 (+/- 0.7) | 10.4 (+/- 0.7) | 9.4 (+/- 0.6) | 9.7 (+/- 0.5) |
| *Carbamidomethyl (excl. C)* | Mouse Lung | 323 (+/- 13) | 67 (+/- 5) | 68 (+/- 18) | 41 (+/- 9) |

## Lab Protocol

### Sample Preparation by Easy Extraction and Digestion (SPEED)

#### Reagents SPEED

- **Trifluoroacetic Acid (TFA)**, ≥ 99 % **Caution**: TFA is highly corrosive. Handle in a fume hood with appropriate personal protective equipment!!!
- **Neutralization Buffer:** 2M TrisBase in H_2_O (12.1 g/50 mL) (pH should not be adjusted!)
- **Reduction/Alkylation Solution (10x):** 100 mM Tris(2-carboxyethyl)phosphine (TCEP, 29 mg/mL), 400 mM 2-Chloroacetamide (CAA, 37 mg/mL) in H_2_O (Prepare fresh before use. Weighed aliquots of TCEP and CAA can be stored at 4°C)
- **Sample Dilution Buffer:** 10:1 (v/v) mixture of 2M TrisBase and TFA
- **H_2_O**
- **Trypsin Stock Solution:** 1µg/µl Trypsin in 50mM acetic acid (e.g. Gold, Mass Spectrometry Grade, Promega)

### Equipment

#### SPEED

- Pipette Tips *(need to be resistant to TFA, e.g. epT.I.P.S. from Eppendorf)*
- Microcentrifuge Tubes (1.5 mL) *(need to be resistant to TFA, e.g epT.I.P.S. from Eppendorf)*
- ThermoMixer
- Microwave (800 Watt)
- *(only needed for certain sample types)*
- UV-VIS Spectrophotometer
- Disposable UV cuvettes (75 -1500 µL)

## Protocol

1. Lyse **cells/tissue** in **TFA** (sample : TFA ~ 1:4 - 1:8 (v/v)) for 2 – 10 min at RT Sample should be mixed occasionally. Usually lysis of cells takes 2-3 min and lysis of tissue up to 10 min. Proteins are stable in TFA for at least 1h (no longer incubation times were tested). Lysis is complete when all cells/tissue are solubilized and DNA is fully degraded (viscosity is water-like).
  1. Gram-positive bacteria require microwave irradiation for 10 s at 800 Watt for efficient lysis.
  2. For safety reasons microcentrifuge tubes are sealed in a 50 mL Falcon tube to prevent TFA leakage if microcentrifuge tube would be damaged in the microwave oven.
  3. Samples should be neutralized instantly after microwave irradiation.
2. Add **Neutralization Buffer** (10 x volume of TFA (step 1)) pH should be ~ 8-9. Samples heat up (exothermic neutralization) and become turbid (proteins precipitate).
3. Add **Reduction/Alkylation Buffer** (1.1 x volume of TFA (step 1))
4. Incubate at 95°C for 5 min Samples can be stored at - 20 or - 80°C for several months
5. Determine protein concentration using **Turbidity Measurement**
  1. Transfer samples to disposable UV cuvettes (dilute with **Sample Dilution Buffer** if necessary)
  2. Measure absorption at 360 nm using a UV-VIS Spectrophotometer
  3. Calculate protein concentration (1AU = 0.79 µg/µL)
  4. Samples can be recovered for further preparation
6. Adjust all samples to 1/0.25/0.05 µg/µL protein concentration using **Sample Dilution Buffer** Use highest possible protein concentration
7. Dilute samples 1:5 with **H_2_O**
8. Add **Trypsin** Use an enzyme to protein ratio (wt:wt) of 1:100 (1 µg/µL), 1:50 (0.25 µg/µL), 1:20 (0.05 µg/µL) depending on 6.
9. Incubate at 37°C and 600 rpm for 20 h in a ThermoMixer Samples can be stored at - 20 or - 80°C for several months
10. Add **TFA** to the samples to a final concentration of 2 % Check pH to be ~ 2
11. Desalt peptides using appropriate protocols, e.g. according to Rappsilber et al. [1]

## Desalting of Peptides (C18-StageTips) [1]

1. Desalt samples with C18-StageTips at 8000 x g for 1-3 min per step Check after each step that all liquid has passed the tip

| Sorbent | X x 1 $\mu\text{m}^3$ C<sub>18</sub> |
| --- | --- |
| **Sample load** | 7 $\mu\text{g}/\mu\text{m}^3$ (14-Gauge: 1 $\mu\text{m}^3$/Plug) |
| **1. Activation** | 100 $\mu\text{l}$ MeOH |
| **2. Equilibration** | 100 $\mu\text{l}$ 0.2 % (v/v) TFA |
| **3. Load** | Sample (check pH to be ~ 2) |
| **4. Wash** | 3 x 200 $\mu\text{l}$ 0.2 % (v/v) TFA |
| **5. Elution** | 100 $\mu\text{l}$ 0.1 % (v/v) FA, 60 % (v/v) ACN |

1. Dry down samples in a SpeedVac Samples can be stored at -20 or -70°C for several weeks
2. Resuspend samples in 20 µL 0.1 % FA

